# Effects of dipicolinic acid on *Bacillus anthracis* spore biology and cytotoxicity

**DOI:** 10.1101/2025.10.10.681759

**Authors:** Chandler Hassan-Casarez, Tiffany V. Mata, Ernesto Abel-Santos

## Abstract

*Bacillus anthracis* is a gram-positive spore-forming bacterium that causes lethal inhalation anthrax. The use of *B. anthracis* as a bioweapon is predicated in its ability to form dormant and resistant infective spores that can be used as agents. *B. anthracis* spores contain large concentrations of dipicolinic acid complexed with cations, especially calcium (Ca-DPA). Following phagocytosis by alveolar macrophages, spores germinate inside the phagolysosome and excrete the Ca-DPA depot into the phagosomal space. In this study, we assessed the effects of DPA on *B. anthracis* spore biology and cytotoxicity. We generated *B. anthracis* mutants with defects in DPA synthesis (*ΔspoVFA, ΔspoVFB, ΔspoVFAB*) or transport (*ΔspoVV*). To increase the viability of DPA-less spores, we also constructed double mutants (*ΔsleBΔspoVFA, ΔsleBΔspoVFB, ΔsleBΔspoVFAB* and *ΔsleBΔspoVV*) by deleting the cortex lytic *sleB* gene. We found that single- and double-mutant DPA-less spores were profoundly compromised in dormancy, viability, germination, and heat resistance. Contrary to expectations, each DPA synthesis mutant exhibited distinct viability and resistance phenotypes. Even with compromised stability, DPA-less *B. anthracis* spores, with the exception of the Δ*sleB*Δ*spoVV* double mutant, were as cytotoxic as wild-type spores. In summary, DPA is required to sustain normal *B. anthracis* spore biology but is not required for macrophage-targeted virulence. Furthermore, the SpoVV transporter and SleB lytic protein seem to have redundant roles in anthrax spore cytotoxicity beyond DPA accumulation.

**Importance:** *Bacillus anthracis* causes deadly pulmonary anthrax and has been used as a weapon for bioterrorism. *B. anthracis* form spores that germinate and establish infection. During germination, *B. anthracis* spores release large amounts of calcium complexed with dipicolinic acid (DPA). In this study, we deleted *the B. anthracis* genes that are required to synthesize and transport DPA into spores. We found that *B. anthracis* DPA-less mutant spores exhibited differential biological effects that were DPA independent. Furthermore, we found that DPA was not required for anthrax cytotoxicity. Finally, we found that proteins involved in DPA synthesis, transport, and cortex lysis have biological and virulence functions that extend beyond DPA accumulation.

## Background

*Bacillus anthracis* is a gram-positive, spore-forming, soil bacterium (1, 2) that has been classified as a Tier 1 select agent used by the CDC (3). Indeed, letters containing *B. anthracis* spores were used as a bioterrorism weapon in 2001 (4, 5).

Similar to other Firmicutes, *B. anthracis* initiates sporulation in response to nutrient limitations (6). During this process, a single bacterium produces a metabolically dormant spore, which is highly resistant to chemical and physical insults (6).

The remarkable dormancy and resistance of spores can be attributed to their unique structures (7). The mineralization of the spore core is one of the most notable protective features (8). Indeed, the low water content and high concentration of cations, especially calcium (Ca^+2^) chelated with dipicolinic acid (Ca-DPA), creates a quasi-crystalline environment that protects biopolymers in the dormant spore. Other important cations, such as magnesium (Mg^+2^), potassium (K^+^), and sodium (Na^+^), also accumulate inside the spore core but at lower concentrations (9).

DPA is synthesized in the mother cell during late sporulation and then transported into the developing spore (10). In *B. subtilis,* the dipicolinate synthase complex, encoded by the *spoVFAB* operon, is produced in the mother cell. Dipicolinate synthase catalyzes the last step of DPA biosynthesis by converting *L*-2,3-dihydrodipicolinate, an intermediate of the lysine biosynthetic pathway, into DPA (10).

A two-step transport pathway has been proposed for the accumulation of DPA inside spores. First, the SpoVV protein, produced in the mother cell, translocates DPA from the mother cell across the outer membrane of the developing spore and into the intermembrane space (11). Proteins from the SpoVA complex produced by developing spores import DPA from the intermembrane space into the spore core (11).

In *B. subtilis*, DPA-less spores are generated by the deletion of the entire *spoVFAB* operon. However, these spores were unstable and either lysed during purification or germinated spontaneously (12, 13). A compensatory deletion of *sleB*, a gene encoding a cortex lytic enzyme, stabilizes *spoVFAB* mutant spores (8). In contrast, *ΔspoVFA, ΔspoVFB,* and *ΔspoVV* mutant spores were isolated without stabilizing mutations (11). These observations suggest that, at least in *B. subtilis*, DPA is required to maintain spore dormancy (13).

Although *B. anthracis* spores are dormant and cannot produce toxins, they serve as infectious anthrax agents (2). When spores sense the restoration of favorable growth conditions, they germinate to resume the vegetative cell cycle (14, 15). Germination is initiated by the recognition of intracellular metabolites, which results in the release of DPA-cation complexes, spore rehydration, and the loss of resistance.

In inhalation anthrax, *B. anthracis* spores are engulfed by resident lung macrophages and transported to the lymph nodes (16). During transit, spores germinate in the phagolysosome while concomitantly releasing DPA and cations inside the infected macrophages (17). Shortly thereafter, newly germinated *B. anthracis* cells begin to secrete toxins that eventually kill the macrophages. Continuous bacterial proliferation and toxin secretion lead to the eventual death of the infected host (18).

DPA is an amphipathic molecule, and the pyridine nitrogen has a pKa close to the pH inside the phagolysosome (19, 20). Thus, during infection, the pyridine nitrogen of DPA could act as a buffer to modulate phagolysosome acidification, providing newly germinated *B. anthracis* cells with sufficient time to produce toxins that lead to macrophage death. Alternatively, spore-released DPA could sequester divalent cations away from macrophages, reducing their bioavailability for host cellular functions.

In this study, we assessed the effects of DPA on *B. anthracis* spore viability, germination, and cytotoxicity of *B. anthracis* spores. We found that deletion of the A-subunit of the DPA synthase complex (but not the B-subunit) leads to overaccumulation of divalent cations inside mutant spores. Increased cation accumulation is lost when the complementary *sleB* gene is deleted. Furthermore, all DPA-less spores were less viable than the wild-type spores. DPA-less spores were more heat-sensitive when the compensating *ΔsleB* mutation was introduced. Notwithstanding their compromised germination and resistance profiles, DPA-less spores, with the exception of Δ*sleB*Δ*spoVV* double mutants, were as cytotoxic to macrophages as wild-type spores.

## Materials and Methods

### Bacterial strain manipulation and growth conditions

*Bacillus anthracis* Sterne 34F2 strain was a generous gift from Dr. Arturo Casadevall (Albert Einstein College of Medicine) and was used to create all DPA-less mutants. *B. anthracis* cells were routinely grown on nutrient broth and agar plates. The allelic exchange plasmids pRP1028 and pRP1099 were generously donated by Dr. Roger Plaut of the United States Food and Drug Administration (USFDA). Allelic exchange procedures were performed on brain heart infusion (BHI) agar and broth supplemented with antibiotics as appropriate.

*E. coli* DH5⍺ was used for subcloning and plasmid construction. *E. coli* CA434 (*E. coli* HB101 harboring pRK24) was used for conjugation with *B. anthracis* cells. *E. coli* cells were cultured in Luria-Bertani (LB) agar and broth supplemented with antibiotics as appropriate. *E. coli* and *B. anthracis* were routinely cultured at 37 °C with shaking at 225 rpm.

All primers were obtained from Integrated DNA Technologies (Coralville, IA, USA). The Wizard Genomic DNA Purification Kit, Benchtop 1 kb, and 100 bp DNA ladders were purchased from Promega Corporation (Madison, WI, USA). The Phusion Hot Start II DNA Polymerase and Zero Blunt TOPO PCR Cloning Kit were purchased from Thermo Fisher Scientific (Waltham, MA, USA). The QIAquick PCR Purification Kit and the Plasmid Mini Kit were purchased from Qiagen (Hilden, Germany). *BamHI* and *BsaI* were purchased from New England Biolabs (Ipswich, MA, USA). Antibiotics were purchased from Sigma-Aldrich (Burlington, MA, USA).

### Construction of DPA-less B. anthracis strains

*B. anthracis* strains lacking DPA were generated following previously published procedures (21). Briefly, the upstream and downstream homology regions for each gene were amplified using appropriate primer sets **(Table S1)**. The resulting upstream and downstream homologous region amplicons were purified and fused by overlap extension PCR (SOE-PCR).

The SOE products were ligated into the TOPO-Zero Blunt vector and transformed into chemically competent *E. coli* DH5α cells. Transformed cells were selected on Luria-Bertani (LB) agar supplemented with 50 µg/ml kanamycin. The resulting pTOPO-SOE plasmids (**Table S2**) were independently purified and digested with *BamHI* and *BsaI*. Each excised SOE construct was ligated into *BamHI*/*BsaI* digested pRP1028 and transformed into chemically competent *E. coli* DH5α. Transformed cells were selected on Luria-Bertani (LB) agar supplemented with 50 µg/ml kanamycin. The resulting pRP1028 plasmids harboring each deletion construct (**Table S2**) were purified and transformed into *E. coli* CA434.

*E. coli* CA434 cultures harboring individual pRP1028 knockout constructs were grown individually for 6 h at 37 °C in LB broth supplemented with 100 μg/ml spectinomycin and 100 μg/ml ampicillin with agitation. The cultures were then pelleted at 2,500 ×g for 5 min and the supernatant was discarded. The pellet was gently resuspended in 1 ml of *B. anthracis* Sterne 34F2, which was grown for 5 h in BHI broth at 37 °C with agitation. The *E. coli/B. anthracis* conjugation mating mixtures were then spotted (7 × 100 µL) onto BHI (without antibiotics) and incubated at 30 °C for 16 h. The conjugation was terminated by harvesting the mating mixture in 1 ml of phosphate-buffered saline (PBS). A 100 µl aliquot of the *E.coli*/*B. anthracis* cell suspension was plated onto BHI supplemented with 250 μg/ml spectinomycin and 110 μg/ml polymyxin B. Plates were incubated for two days at 30 °C.

After two days, the plates were viewed under green light (520 nm) using a Typhoon 9410 Variable Mode Imager to screen for single integrants. Fluorescent colonies were passed onto fresh BHI plates supplemented with 250 μg/ml spectinomycin and 110 μg/ml polymyxin B. The plates were incubated overnight at 37 °C. The increased temperature was non-permissive for plasmid replication and was selected against individuals harboring non-integrated pRP1028. The temperature-dependent selection process was repeated thrice.

To stimulate the second recombination event, *B. anthracis* integrants were conjugated with *E. coli* CA434 harboring pRP1099, using the same procedure described above. After terminating conjugation, *E. coli*/*B. anthracis* mixture was plated onto BHI medium supplemented with 20 μg/ml kanamycin and 110 μg/ml polymyxin B and incubated overnight at 37 °C. Single colonies from the overnight incubation were picked and streaked onto fresh BHI supplemented with 20 μg/ml kanamycin and 110 μg/ml polymyxin B. Plates were incubated overnight at 37 °C.

Following overnight incubation, the plates were imaged under green light (520 nm) to screen for the loss of pRP1028. Non-fluorescent colonies were picked and patched onto BHI supplemented with 250 μg/ml spectinomycin and 110 μg/ml polymyxin B. Colonies were also replicated and patched onto unsupplemented BHI medium. All plates were incubated overnight at 37 °C. The next day, spectinomycin-sensitive strains were PCR-screened for deletions using the appropriate primers, passed onto BHI, and incubated overnight at 37 °C.

Following this overnight incubation, the plates were viewed under blue light (455 nm) to screen for the loss of pRP1099. Non-fluorescent colonies were patched onto BHI medium supplemented with 20 μg/ml kanamycin and 110 μg/ml polymyxin B. Colonies were also replicated and patched onto unsupplemented BHI medium. All plates were incubated overnight at 37 °C. The next day, glycerol stocks of the kanamycin-sensitive deletion strains **(Fig. S1)** were prepared and stored at -80 °C.

Double mutants were created by conjugating the corresponding single mutants with *E. coli* carrying pRP1028 containing deletions regions of *sleB* (**Table S2**). Double mutants were selected using the procedure described above.

To confirm single and double deletions, genomic DNA from wild-type *B. anthracis* and putative *B. anthracis* mutants were used as PCR targets. Regions flanking the *spoVFAB* operon from *B. anthracis* Sterne 34F2 were amplified using primer pairs 1020 and 1021 to generate 3500 bp amplicons from *B. anthracis* wild-type, 2600 bp amplicons from *B. anthracis ΔspoVFA*, 2900 bp amplicon from *B. anthracis ΔspoVFB,* and 2000 bp amplicon from *B. anthracis ΔspoVFAB* cells. Regions flanking the *spoVV* gene were amplified using primer pairs 1022 and 1023 to generate 1555 bp amplicons from *B. anthracis* wild-type and 343 bp amplicons from *B. anthracis ΔspoVV* . Lastly, regions flanking *sleB* were amplified using primer pairs 1024 and 1025 to generate a 2235 bp amplicon from *B. anthracis* wild-type and 603 bp amplicons from *B. anthracis ΔsleB* cells (**Fig. S2**).

Each confirmation amplicon was then purified, and the DNA was quantified using Nanodrop and sequenced by Genewiz to confirm genomic deletions. The *spoVFAB* amplicon was sequenced using the primer pairs 1026 and 1027. The *spoVV* amplicon was sequenced using the primer pairs 1028 and 1029. The *sleB* amplicon was sequenced using the primer pairs 1030 and 1031 **(Table S3)**. Sequencing results were translated and aligned with the *Bacillus anthracis* Sterne 34F2 genome using benchling to confirm genomic deletions.

### B. anthracis spore preparation and purification

Frozen stocks of wild-type *B. anthracis* cells were streaked onto nutrient agar plates and incubated overnight at 37 °C to yield single colonies. Liquid cultures of *B. anthracis* were prepared by inoculating nutrient broth with a single *B. anthracis* colony and grown at 37 °C with agitation for 3-4 hours. Aliquots (200 μl) from the liquid *B. anthracis* culture were spread onto sporulation plates (nutrient agar supplemented with 10% KCl, 1.2% MgSO_4_, 1 M Ca(NO_3_)_2_, 10 mM MnCl_2_, and 1 mM FeSO_4_) and incubated at 37 °C for six days to allow for sporulation. The resulting bacterial lawns were harvested by scraping in ice-cold deionized water (DI-H_2_O). The spores were pelleted by centrifugation at 8,500 xg for 5 min at 4 °C. The supernatants were decanted, and the spore pellets were resuspended in DI-H_2_O. This process was repeated thrice. Following the third wash with DI-H_2_O, the spore pellet was resuspended in 20% HistoDenz solution, layered on top of a 50% Histodenz solution, and centrifuged at 11,500 ×g for 35 min (no brake). After discarding the Histodenz solution, the resulting spore pellet was washed three times by repeated centrifugation (8,500 xg, 5 min, 4 °C) and resuspension in DI-H_2_O. The spores were stored in DI-H_2_O at 4 °C, until needed. Spore preparations were >95% pure, as determined by Schaeffer–Fulton staining and by microscopic observation under phase-contrast conditions.

*B. anthracis* DPA-less cells were sporulated, as described above. Because of the fragility of DPA-less spores, mutant spores were purified by repeated washes and resuspensions in DI-H_2_O instead of gradient centrifugation. Following the third wash with DI-H_2_O after harvesting, 5 ml of DPA-free spores were resuspended in 45 ml of DI-H_2_O and stored at 4 °C for at least 24 h. During this period, debris was separated from the intact spores and floated on top of the supernatant. The supernatant was then removed and the spore pellet was resuspended in 45 ml of DI-H_2_O. This process was repeated until no debris floated in the supernatant. Spore preparations were >95% pure as determined by microscopic observation (22).

### Microscopy observation of B. anthracis spores

Aliquots of *B. anthracis* spores were placed on a microscope slide and sandwiched with a cover glass. Spores were microscopically observed on a Motic BA310 microscope with a 40X objective in the phase contrast mode. Images were acquired with a Moticam A16 camera.

### B. anthracis spore cation content

Metal ion content was determined via inductively coupled plasma atomic emission spectroscopy (ICP-AES) using a Shimadzu ICP-9000 instrument. Aliquots of purified spores were resuspended in 5 ml of DI-H_2_O to OD_580_ = 1.0. The spores were then diluted with 15 ml of 2% HNO_3._ Spore suspensions were incubated overnight at 90 °C. The dissolved spore suspension was cooled to room temperature and centrifuged (11,500 xg, 5 min) to pellet debris. The supernatant was then transferred to a fresh polypropylene tube and autoclaved (121°C for 20 min). After cooling to room temperature, the solution was filtered through a 0.2 μm PES filter into a fresh 15 ml conical tube. Prior to analysis, filtrates were tested for sterility by plating 10 µl aliquots onto nutrient agar and incubating at 37 °C for 12 h. Cation concentrations were determined by interpolation of standard curves using the ICP Solution software.

### Fast germination of B. anthracis spores

Spore germination was monitored spectrophotometrically using a LabSystems iEMS 96-well plate reader equipped with a 580 nm cut-off filter (OD_580_). Germination experiments were performed in clear flat-bottomed 96-well plates. Aliquots of purified spores were individually resuspended in germination buffer (50 mM Tris-HCl, 10mM NaCl, pH 7.5) and diluted to OD_580_ = 1.0. The optical density of the spore suspensions (160 μL) was measured for 15 min to monitor auto-germination. Germination reactions were then started by adding 20 μL of a 0.4 mM L-alanine solution and 20 μL of a 0.4 mM inosine solution to each spore suspensions (200 μL/well final volume). The optical densities were measured every 1-minute for 60 min. The relative OD_580_ was calculated by dividing each OD_580_ measurement by the original OD_580_ obtained at time zero [Relative OD_580_ = OD_580_(t)/OD_580_(t_0_)]. Experiments were performed in triplicates (technical replicates) with at least three different spore preparations (biological replicates). The samples were not heat-activated prior to the experiments because of the susceptibility of DPA-free spores to wet heat.

### Detection of DPA content in B. anthracis spores

Total DPA content was measured by monitoring the fluorescence intensity of the Tb^+3^-DPA complex in a Tecan Infinite M200 plate reader at an excitation wavelength of 270 nm and an emission wavelength of 545 nm. Fluorescence assays were performed in black 96-well plates with flat and clear bottom. Aliquots of purified spores were prepared as described above and diluted to OD_580_ = 1.0 in germination buffer. Aliquots of spores were lysed by heating at 95 °C for two hours. Each reaction contained 30 μL of lysed spore suspension, 35 μL of a 250 μM TbPV (250 μM TbCl_3_ and 250 μM pyrocatechol violet) solution, and 85 μL of germination buffer (150 μL/well final volume). The total DPA concentration was determined by interpolating the fluorescence signals into a Tb-DPA calibration curve.

The lack of DPA release during fast germination of *B. anthracis* mutant spores was further confirmed by LC-MS. Wild-type and DPA-less mutant spores germinated in the presence of l-alanine and inosine as described above. Supernatants were then filtered through a 0.2 µm PES filter. The samples were streaked onto BHI plates and incubated for 48 h to ensure sterility. Sample analysis was performed at The Metabolomics Innovation Centre (TMIC; Edmonton, Alberta, CA, USA). Samples were analyzed using a Shimadzu Nexera XR UHPLC interfaced with a Sciex 6600 TripleToF MS operating in the positive ion mode. The MS data were analyzed and quantified using Sciex PeakView software.

### B. anthracis delayed spore germination, viability, and heat resistance

Spore germination, viability, and heat resistance were assessed by enumeration of colony forming units (CFUs). Samples of purified spores were prepared as described above and diluted to OD_580_ = 1.0 in DI-H_2_O. Each spore sample was divided into three aliquots. One aliquot was maintained at room temperature (to test for delayed spore germination and viability). The second aliquot was heated at 68 °C for 30 minutes (to test spore resistance to moderate heat). The third aliquot was heat-treated at 90 °C for 5 minutes (to test spore resistance to high heat). All heated and nonheated spore aliquots were serially diluted, plated on nutrient agar, and incubated overnight at 37 °C. CFUs were enumerated on plates showing non-confluent colonies.

### Mammalian cell line and growth conditions

The murine monocyte/macrophage RAW 264.7 cell line was purchased from America Type Culture Collection (Manassas, VA, USA). RAW 264.7, cells were grown routinely in Dulbecco Modified Eagle’s medium (DMEM) with 4.5 g/l D-glucose and sodium pyruvate, without L-glutamine. DMEM was supplemented with 10% fetal bovine serum, 1% GlutaMAX, and 1% PenStrep (growth medium). Cells were grown in 75 cm^2^ cell culture flasks in 10 ml of medium. The flasks were incubated at 37 °C in a humidified atmosphere containing 5% CO_2_.

When cells reached 80-90% confluency, they were sub-cultured by incubating at room temperature for 2 min with 10 ml of PBS supplemented with 5 mM EDTA. Cells were detached from the flask by scraping. Cell suspensions were collected and centrifuged at 500 xg for 5 min. The cell pellets were resuspended in growth medium. Viable cell counts were determined by trypan blue exclusion using a hemocytometer. Subcultures were seeded in new flasks at a final concentration of 300,000 viable cells/ml. Subcultures were routinely performed every four days. After the 20^th^ subculture, the health of cell lines deteriorated, and was discarded. A new cell culture was initiated using frozen liquid nitrogen stocks.

### Macrophage cytotoxicity assay by B. anthracis spores

RAW264.7 cells were seeded into black 96-well plates with clear flat bottoms using 200 µl of 50,000 cells/ml suspension. The cells were incubated for 36 h at 37 °C with 5% CO_2_. During this period, the cells grew to near confluence (80,000 cells/well).

After 36 h, the growth medium was removed and the cells were washed three times with PBS. Cells were then incubated at 37 °C with 5% CO_2_ for 1 h in DMEM supplemented with 1% GlutaMAX. *B. anthracis* spores were added at the appropriate multiplicity of infection (MOI) in volumes of 100 μL. Immediately following spore challenge, cell cultures were centrifuged at 500 xg for 5 min to ensure contact between macrophages and spores. The plates were incubated for 90 min at 37 °C in an atmosphere of 5% CO_2_.

Following the spore challenge, the cells were washed three times with PBS to remove non-phagocytized extracellular spores. After washing, the infected cells were incubated with DMEM supplemented with 10% horse serum, 1% GlutaMAX, and 1.25 μg/ml gentamycin until measurements were recorded.

To determine anthrax-mediated macrophage killing, propidium iodide (PI) was added to each well at a final concentration of 20 μM. Fluorescence intensities were measured using a Tecan Infinite M200 plate reader at an excitation wavelength of 535 nm, emission wavelength of 617 nm, and gain of 100. Prior to each measurement, the plates were shaken for 10 seconds. Fluorescence readings were taken every hour, beginning at 2 h post-infection. The data were obtained in relative fluorescence units (RFU) and converted to relative intensity (RI) using the following formula:

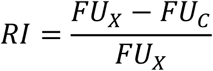

Where FU_x_ is the specific fluorescence signal of each condition, and FU_c_ is the fluorescence value of the uninfected control. This conversion was corrected for variations in the background fluorescence from one experiment to another.

RI was then converted to percent macrophage killing defined as:

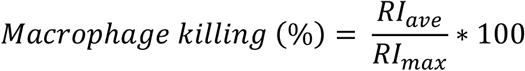

Where RI_ave_ corresponds to the average RI of the infected cells and RI_max_ corresponds to the highest average RI of the positive control (macrophages treated with 1% tritonX100).

### Statistical analysis

Data were visualized using Microsoft Excel LTSC MSO (16.0.14334.20090) 64-bit. All measurements were made with at least three different spore preparations (biological replicates) and at least three separate measurements per assay (technical replicates). The means and standard deviations were calculated from at least nine data points using Microsoft Excel.

One-way single-factor analysis of variance (ANOVA) or one-tailed t-test was performed, as appropriate, using R Statistical Software (v. 4.4.3)(23). Statistical differences between spore cation concentrations, spore viability, and spore cytotoxicity were determined using ANOVA to assess differences among different groups. Significant main-effect ANOVAs (*p*<0.05) were analyzed using Scheffe’s correction for *post hoc* comparisons between all groups. Statistically significant differences between groups were defined as *p*<0.05. Statistical differences in heat resistance between the heat-treated and untreated spores were determined using a one-tailed unpaired t-test.

## Results

### Generation of DPA-less B. anthracis spores

DPA-less spores were created by deletion of *B. anthracis spoVFA*, *spoVFB*, *spoVFAB* or *spoVV* genes. Single mutants were subjected to a second round of gene deletion targeting the cortex-lytic enzyme SleB to yield *B. anthracis ΔsleBΔspoVFA*, *ΔsleBΔspoVFB, ΔsleBΔspoVFAB,* and *ΔsleBΔspoVV* double mutants. The identity of all mutants was confirmed by PCR (**Fig. S2)** and subsequent sequencing. As expected, spores from all *B. anthracis* mutants failed to accumulate DPA (**Table S4**). To confirm the lack of DPA in the mutant spores, all spores were screened for DPA release during germination using mass spectrometry. DPA was detected in the media after the germination of wild-type *B. anthracis* spores **(Fig. S3A)**. As expected, no DPA was released into the germination media of DPA-less *B. anthracis* mutants spores **(Fig. S3B)**.

### Microscopy of DPA-less B. anthracis spores

Phase contrast microscopy of wild-type spores showed a homogenous suspension of phase-bright particles (**Fig. S4A)** and stained green with Schaeffer–Fulton staining. In contrast, DPA-less spores revealed populations of phase-bright (white-capped spores, bright halo), phase-gray (dark-capped spores, dim halo), and phase-dark spores (no cap, no halo) **(Fig. S4B-S4C)**. Similarly, DPA-less spores showed green, red, and brown particles when stained with Schaeffer–Fulton stain.

### Cation content of DPA-less B. anthracis spores

The Δ*spoVFA* and Δ*spoVFAB* mutant spores showed altered mineralization with significantly higher levels of Ca^+2^, Mg^+2^, and, to a lesser extent, Na^+^ cations, compared to wild-type *B. anthracis* spores. The increased accumulation of cations in Δ*spoVFA* and Δ*spoVFAB* mutant spores was reversed to wild-type levels when *sleB* was also deleted (**Figs. 1A-1E**).

**Figure 1.**
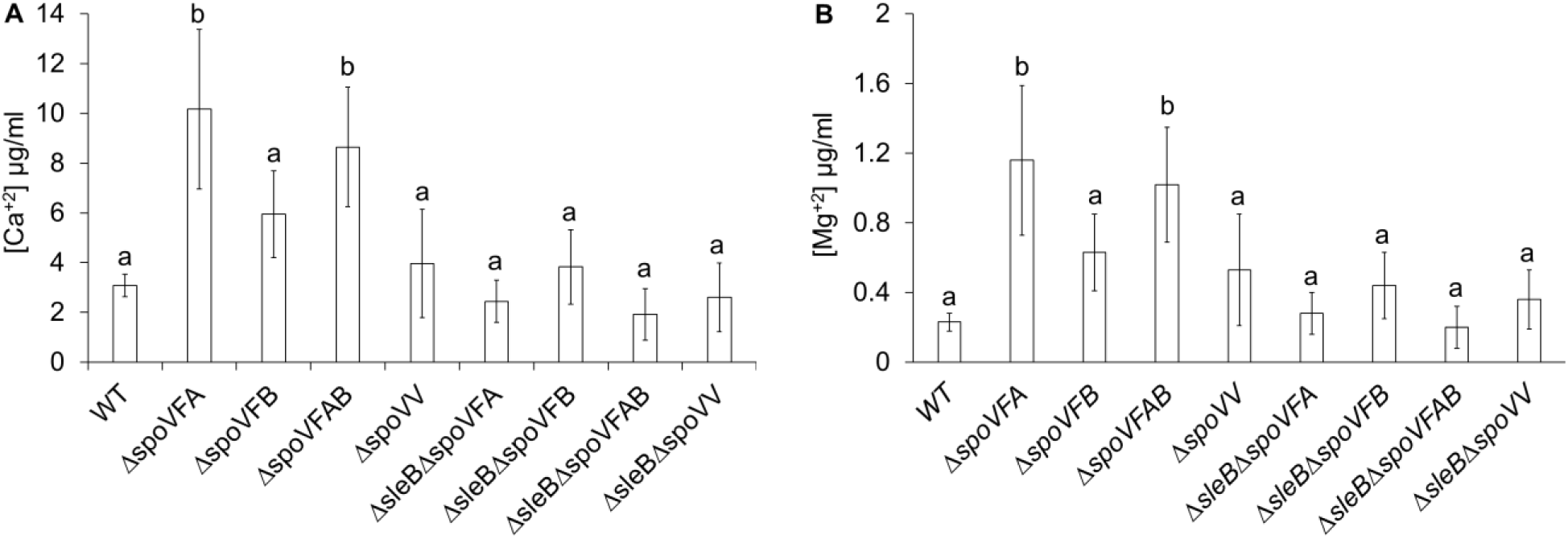
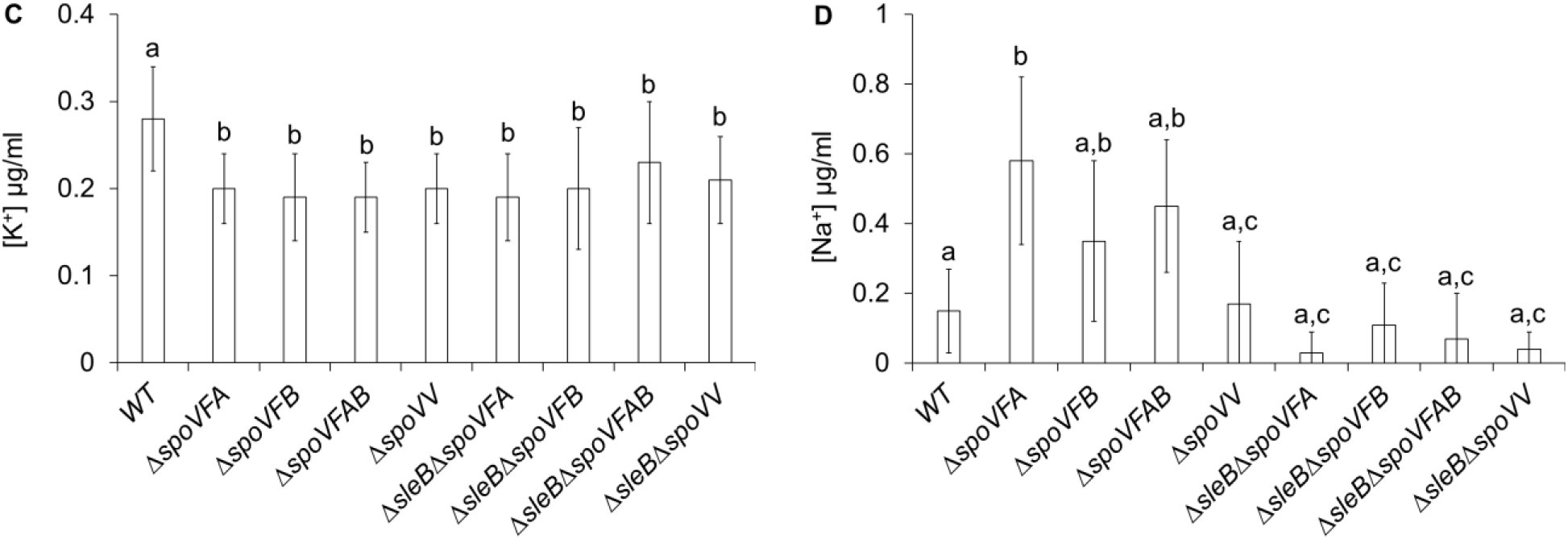
Cation content of DPA-less *B. anthracis* spores. Suspensions of *B. anthracis* wild type (WT), Δ*spoVFA,* Δ*spoVFB,* Δ*spoVFAB,* Δ*spoVV,* Δ*sleB*Δ*spoVFA*, Δ*sleB*Δ*spoVFB*, Δ*sleB*Δ*spoVFAB*, and Δ*sleB*Δ*spoVV* spores were individually diluted to OD_580_= 1.0 to a final volume of 5.0 ml and supplemented with 2% HNO_3_. Suspensions were incubated overnight at 90 °C. **(A)** calcium, **(B)** magnesium , **(C)** potassium, and **(D)** sodium concentrations were determined by interpolation of standard curves using ICP Solution software. At least three independent biological replicates were measured. Each biological replicate was measured three times as technical replicates. Data are presented as mean ± SD. Single-factor ANOVA was performed to assess differences in cation content among different spore mutants. Significant main-effect ANOVAs (*p*<0.05) were analyzed using Scheffe’s correction for *post hoc* comparisons between all groups. Columns that are labeled with different lowercase letters indicate groups that are statistically different (*p*<0.05 between groups).

### Fast germination of DPA-less B. anthracis spores

Wild-type spores showed a rapid drop in optical density when germinated in the presence of inosine and L-alanine **(Fig. 2)**. In contrast, there were no changes in the optical density for any DPA-less *B. anthracis* mutant spores, even after 1 h of incubation. None of the spore suspensions showed auto-germination prior to incubation with inosine or L-alanine.

**Figure 2.**
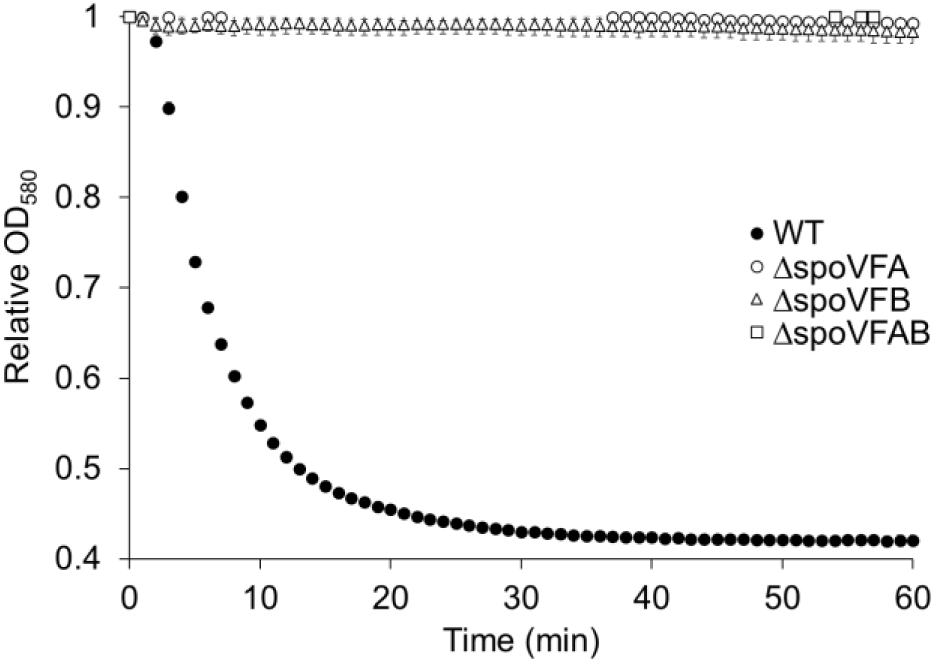
Fast germination of DPA-less *B. anthracis* spores. Wild type and DPA-less mutant *B. anthracis* spores were treated with inosine and L-alanine at 0.4 mM final concentrations. Optical density data were collected at 580 nm at 1-minute intervals for 60 minutes. Data represent averages from three independent measurements and error bars represent standard deviations. In the presence of inosine and L-alanine, wild-type spores (black circles) show fast decrease in optical density that correspond to spore germination. In contrast, *B. anthracis* Δ*spoVFA* (white circles), Δ*spoVFB* (white triangles), and Δ*spoVFAB* (white squares) did not show any decrease in optical density even after 60-minute incubation with inosine and L-alanine. Other DPA-less spores showed similar patterns and were omitted for clarity.

### Viability and heat resistance of DPA-less B. anthracis spores

All DPA-less spores were viable but formed fewer colonies than wild-type spores (**Fig. 3, white columns**). *B. anthracis ΔspoVFB, ΔspoVV, ΔsleBΔspoVFB,* and *ΔsleBΔspoVV* spores lost less than 1-order of magnitude CFUs compared to wild-type spores. On the other hand, *B. anthracis ΔspoVFA, ΔsleBΔspoVFA, ΔspoVFAB* and *ΔsleBΔspoVFAB* spores formed between 1.4 and 3.0 orders of magnitude fewer colonies than wild-type spores.

**Figure 3.**
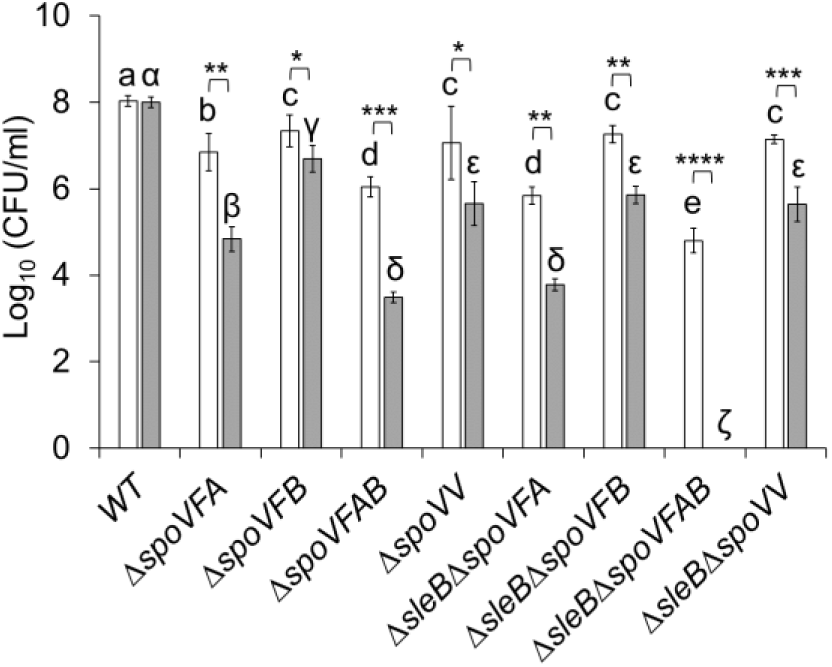
Viability of DPA-less *B. anthracis* spores. Aliquots (OD_580_ = 1.0) of wild-type (WT), *ΔspoVFA, ΔspoVFB, ΔspoVFAB, ΔspoVV, ΔsleBΔspoVFA, ΔsleBΔspoVFB, ΔsleBΔspoVFAB* and *ΔsleBΔspoVV* spores were plated before (white columns) and after (gray columns) incubation for 5 minutes at 68 °C. Spore suspensions were serial diluted, plated on nutrient agar, and incubated for 12 hours at 37 °C. Spore viability was determined by enumeration of colony forming units (CFUs). At least three independent biological replicates were measured. Each biological replicate was measured three times as technical replicates. Data are presented as mean ± SD. Single-factor ANOVA was performed to assess differences in viability among different spore mutants. Significant main-effect ANOVAs (*p*<0.05) were analyzed using Scheffe’s correction for *post hoc* comparisons between all groups. Columns labeled with different lowercase letters indicate groups that are statistically different among untreated spores (white columns, *p*<0.05 between groups). Columns that are labeled with different Greek letters indicate groups that are statistically different among heat-treated spores (gray columns, *p<*0.05 between groups). Statistical differences between matched untreated (white columns) and heat-treated (gray columns) spores were determined by one-tailed unpaired t-test. Asterisks denotes statistically significant differences (*, *p*<0.05; **, *p*<0.01; ***, *p*<0.005; ****, *p*<0.001).

The viability of all DPA-less spores was significantly reduced after heating at moderate temperatures **(Fig. 3, gray columns)**. *B. anthracis ΔspoVFB* spores were the least heat sensitive and formed only 0.7-orders of magnitude less colonies after heating. All other DPA-less mutants lost between 1.3 and 2.6 orders of magnitude of viability after heat treatment. *B. anthracis ΔsleBΔspoVFAB* spores were highly heat-sensitive, and no colonies were recovered after heating. Under high-heat conditions, only wild-type spores were able to form colonies (data not shown).

### Cytotoxicity of DPA-less B. anthracis spores

At higher MOIs, spores with defects in DPA synthesis (Δ*spoVFB* and *ΔsleB*Δ*spoVFB*) showed cytotoxicity similar to that of wild-type spores, with most macrophages dying by three hours post-challenge **(Fig. 4A)**. *B. anthracis* spores with defects in DPA transport (*ΔspoVV* and *ΔsleB*Δ*spoVV*) showed delayed cytotoxicity, with statistically lower macrophage killing at earlier time points. However, at 6-hours post-challenge, macrophage killing by *ΔspoVV* and *ΔsleB*Δ*spoVV* spores was indistinguishable from DPA synthesis mutant spores or wild-type spores.

**Figure 4.**
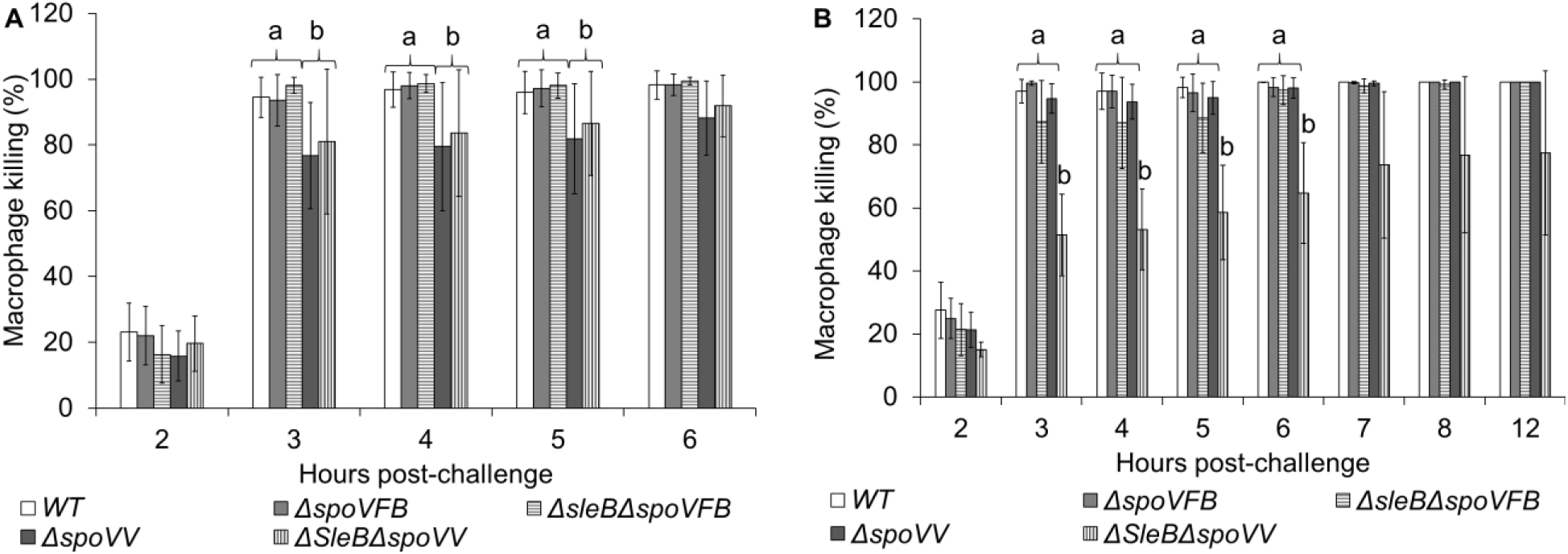
Cytotoxicity of DPA-less *B. anthracis* spores. Macrophage cultures were challenged with either (**A**) 80 MOIs or (**B**) 20 MOIs of *B. anthracis* wild type (WT, white columns), Δ*spoVFB* (light gray columns), Δ*sleB*Δ*spoVFB* (horizontal dashed columns), Δ*spoVV* (dark gray columns),or Δ*sleB*Δ*spoVV* (vertical dashed columns) spores. Cell viability was determined by propidium iodide exclusion at different time points after spore challenge. At least three independent biological replicates were measured. Each biological replicate was measured three times as technical replicates. Data are presented as mean ± SD. Single-factor ANOVA was performed to assess differences in cytotoxicity among different spore mutants. Significant main-effect ANOVAs (*p*<0.05) were analyzed using Scheffe’s correction for *post hoc* comparisons between all groups. Columns labeled with different lowercase letters indicate groups that are statistically different among untreated spores (white columns, *p*<0.05 between groups).

At lower MOIs, *ΔsleB*Δ*spoVV*, but not *ΔspoVV*, mutant spores showed lower cytotoxicity **(Fig. 4B)**. At later time points, cell death by *ΔsleB*Δ*spoVV* spores became heterogeneous. Indeed, some *ΔsleB*Δ*spoVV* samples still showed over 50% macrophage survival, even 12 h post-challenge. In contrast, every other DPA-less mutant caused complete macrophage death within the same timeframe.

## Discussion

In this study, we assessed the role of DPA in *B. anthracis* spore stability, viability, germination, and cytotoxicity. Phase contrast microscopy revealed that all mutant DPA-less *B. anthracis* spores formed populations of bright-, gray-, and dark-phase particles. Schaeffer–Fulton staining further confirmed the presence of at least three different spore populations. This suggests that all mutant spores underwent at least some early germination events (phase gray particles) or late germination events (phase dark particles).

The proportions of phase-bright, phase-gray, and phase-black spores were difficult to quantify accurately. By necessity, phase contrast microscopy must be performed without fixing the spore preparations. Hence, spores randomly moved in and out of the microscope field. However, we estimated that in DPA-less *B. anthracis* spores, the majority of the populations were either phase gray or phase dark.

In *B. anthracis*, inosine and L-alanine trigger a fast germination response that can be monitored in real-time by a decrease in optical density (24, 25). As spores quickly release their mineral deposits and rehydrate, they transition from the bright to the dark phase. Even though all mutant spores still accumulated cations (in some cases at higher levels than wild-type spores), none of the DPA-less *B. anthracis* spores showed a measurable decrease in optical density in response to inosine and L-alanine.

Although mutant spores did not show fast germination, they formed colonies on nutrient agar, albeit at lower levels than wild-type spores. Because colony formation is preceded by germination, these results show that DPA-less *B. anthracis* spores germinate slowly.

The sum of these results suggests that since the majority of DPA-less *B. anthracis* spores start as phase gray/dark particles, the population of fast-germinating (phase-bright) spores might not be sufficient to be detected by optical density. Nevertheless, we suspect that phase-gray (but not phase-black) spores are still viable, accounting for the formation of a reduced but measurable number of colonies. Because phase-gray spores are partially germinated, they would have lost some of their intrinsic resistance, as shown by the decrease in colony formation after heat treatment.

The high cation content of spores is believed to be an important factor contributing to their wet-heat resistance (13, 26). Although most of the DPA-less spores showed cation concentrations that were similar to those of wild-type spores, all mutant spores were significantly less heat-resistant. Interestingly, spores derived from cells that lack the SpoVFA protein (Δ*spoVFA* and Δ*spoVFAB*) showed significantly higher levels of divalent cations but were still heat sensitive. Thus, even high cation concentrations were insufficient to provide heat resistance in the DPA-less spores.

In other *Bacilli* (8), DPA-less spores were unstable unless the cortex-lytic enzyme SleB was deleted (8). However, deletion of the *B. anthracis sleB* gene did not lead to increases in phase brightness, germination, viability, or heat resistance in any of the tested DPA-less spores. In contrast, the deletion of *sleB* caused a significant reduction in the viability of *ΔspoVFA, ΔspoVFB,* and especially *ΔspoVFAB* spores. In contrast, *ΔspoVV* spores were not affected by concurrent deletion of *sleB*.

The lytic activity of sleB is sensitive to the structure of its peptidoglycan substrates (27). Thus, the results from our deletion mutants suggest that compromising DPA synthesis creates spore peptidoglycan structures that are different from those created from impaired DPA transport. We are currently testing this possibilities.

Surprisingly, *B. anthracis* Δ*spoVFAB* spores were orders of magnitude less viable and heat-resistant than Δ*spoVFA* or Δ*spoVFB* spores, since deletion of either subunit stopped DPA production. Furthermore, both the A- and B-subunits are predicted to be globular proteins that are produced by the mother cell and are localized in the cytoplasm. Thus, it is unclear how the removal of individual subunits or the entire SpoVFAB complex differentially alters spore physiology in a DPA-independent manner.

The role of DPA in anthrax virulence has not yet been investigated prior to this study. Following established protocols, we initially attempted to challenge macrophages with MOIs based on the number of *B. anthracis* spores that could form colonies (24, 25). However, because of the low viability of DPA-less spores, attempts to infect macrophages with standardized CFU counts (see **Fig. 3**) resulted in large excesses of unphagocytized particles that could not be removed without damaging the host cells.

To circumvent this problem, we infected macrophages with inoculums standardized by the sum of bright, gray, and dark phase particles, regardless of their viability. To further reduce unphagocytized particles, we tested only the most viable mutant spores (*ΔspoVFB, ΔsleBΔspoVFB, ΔspoVV,* and *ΔsleBΔspoVV*).

Contrary to expectations, wild-type *B. anthracis* spores and most DPA-less spores showed similar cytotoxicity. This suggests that phase-gray and/or phase-dark spores must be able to infect macrophages. Furthermore, these data showed that DPA is not an important determinant of macrophage invasion and death.

In most cases, mutant spore cytotoxicity was not altered after *sleB* deletion. The sole exception was Δ*sleB*Δ*spoVV* spores, which showed lower cytotoxicity at low MOIs than other spores. Because the levels of mineralization of the Δ*spoVV* and Δ*sleB*Δ*spoVV* mutant spores are indistinguishable, the lower cytotoxicity of the Δ*sleB*Δ*spoVV* mutant spores cannot be attributed to either a lack of DPA or ion effects. Our results suggest that the membrane protein SpoVV and lytic enzyme SleB have redundant roles in anthrax spore cytotoxicity that goes beyond DPA transport.

If DPA was the only factor affecting mutant spore physiology, it would be expected that all mutant spores would show similar phenotypes. However, DPA-less spores show diverse cation contents, viability, heat resistance, and cytotoxicity, depending on which genes are deleted. In fact, it seems that SpoVFAB synthase, SpoVV transporter, and SleB lytic protein all uniquely contribute to spore stability and/or cytotoxicity, regardless of their enzymatic activity.

## Acknowledgments

We would like to thank Prof. Arturo Casadevall and Dr. Roger Plaut for their generous contribution to the materials used in this project. We would also like to acknowledge Prof. Vern Hodge *in memoriam* for his help in developing methods for ICP-AES.

## Author contributions

CHC designed and performed the experiments, collected and analyzed the data, wrote the original draft of the manuscript, and edited the manuscript. TVM performed the experiments, collected data, and edited the final manuscript. EAS designed the project and experiments, analyzed the data, acquired funding, and edited the manuscript.

## Data access

All data will be available by request to the corresponding author.

---

**Figure S1.**
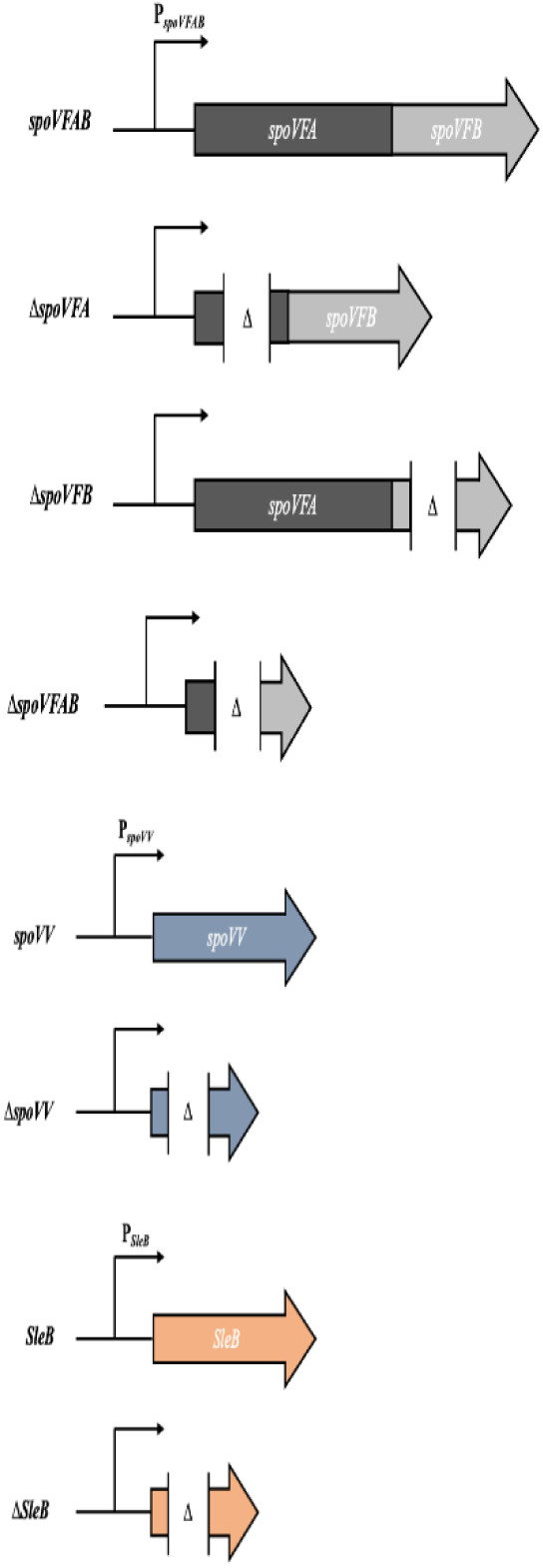
*B. anthracis* gene deletions performed in this study. Deletions of *spoVFA, spoVFB, spoVFAB, spoVV,* and *SleB* in *B. anthracis* Sterne 34F2. Each construct generates a putative non-functional, three amino acid peptide corresponding to the first two amino acids and the last amino acid encoded by the respective knockout.

**Figure S2.**
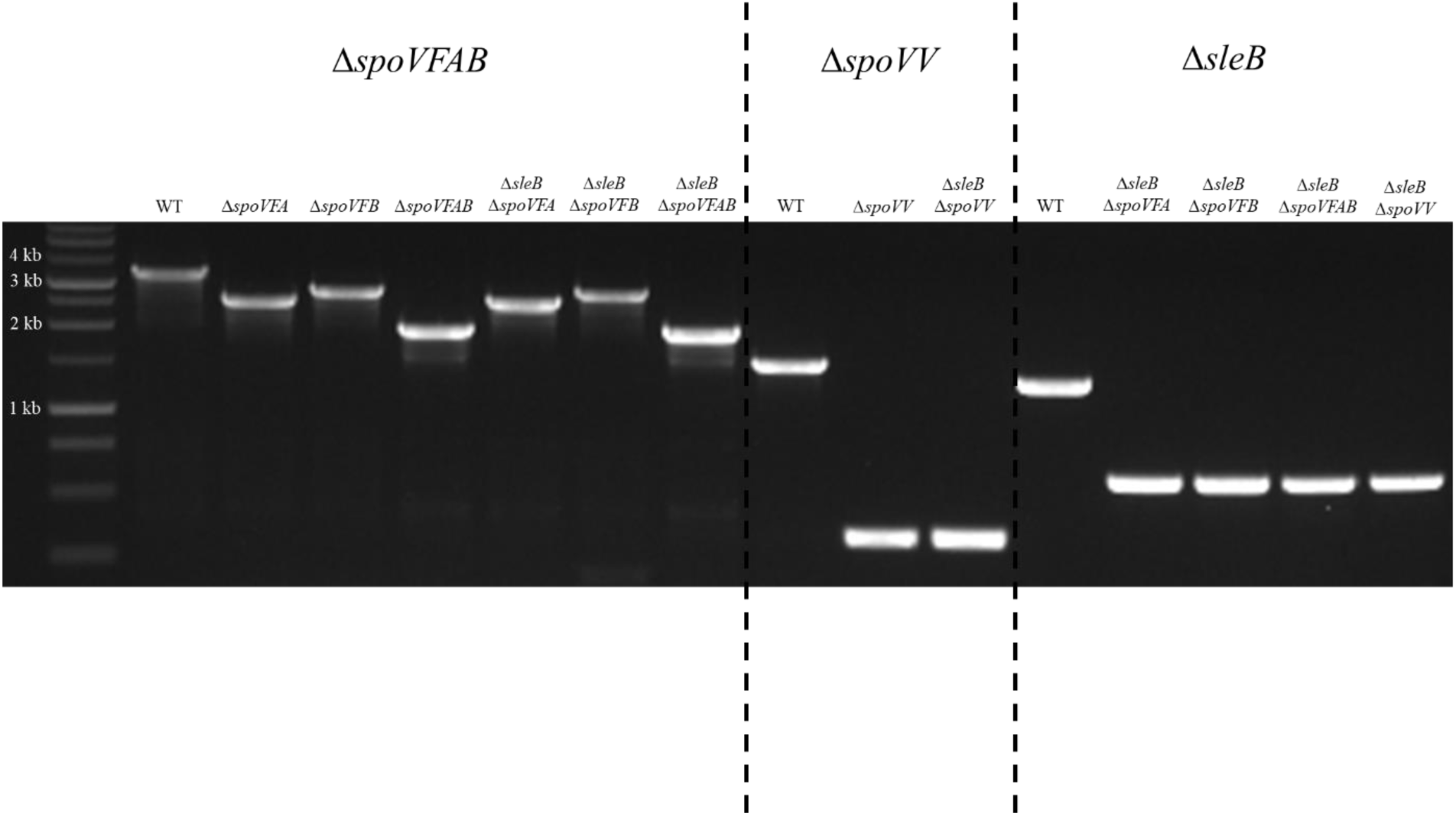
Verification of *spoVF, spoVV*, and *sleB* deletions by colony PCR. In the first panel, mutants were screened for the presence of the *spoVFAB* operon. Wild type *B. anthracis* shows the expected 3500 bp amplicon. *B. anthracis ΔspoVFA* and *ΔsleBΔspoVFA* show a 2600 bp amplicon. *B. anthracis ΔspoVFB* and *ΔsleBΔspoVFB* show a 2900 bp amplicon. *B. anthracis ΔspoVFAB* shows a 2000 bp amplicon. In the second panel, mutants were screened for the presence of the *spoVV* gene. Wild type *B. anthracis* shows the expected 1555 bp amplicon. *B. anthracis ΔspoVV* and *ΔsleBΔspoVV* show a 343 bp amplicon. In the third panel, double mutants were screened for the presence of the *sleB* gene. Wild type *B. anthracis* shows the expected 1338 bp amplicon. All double mutants *B. anthracis* show a *ΔsleB* 603 bp amplicon. A BenchTop 1 kb DNA ladder was used to interpolate amplicon sizes.

**Figure S3.**
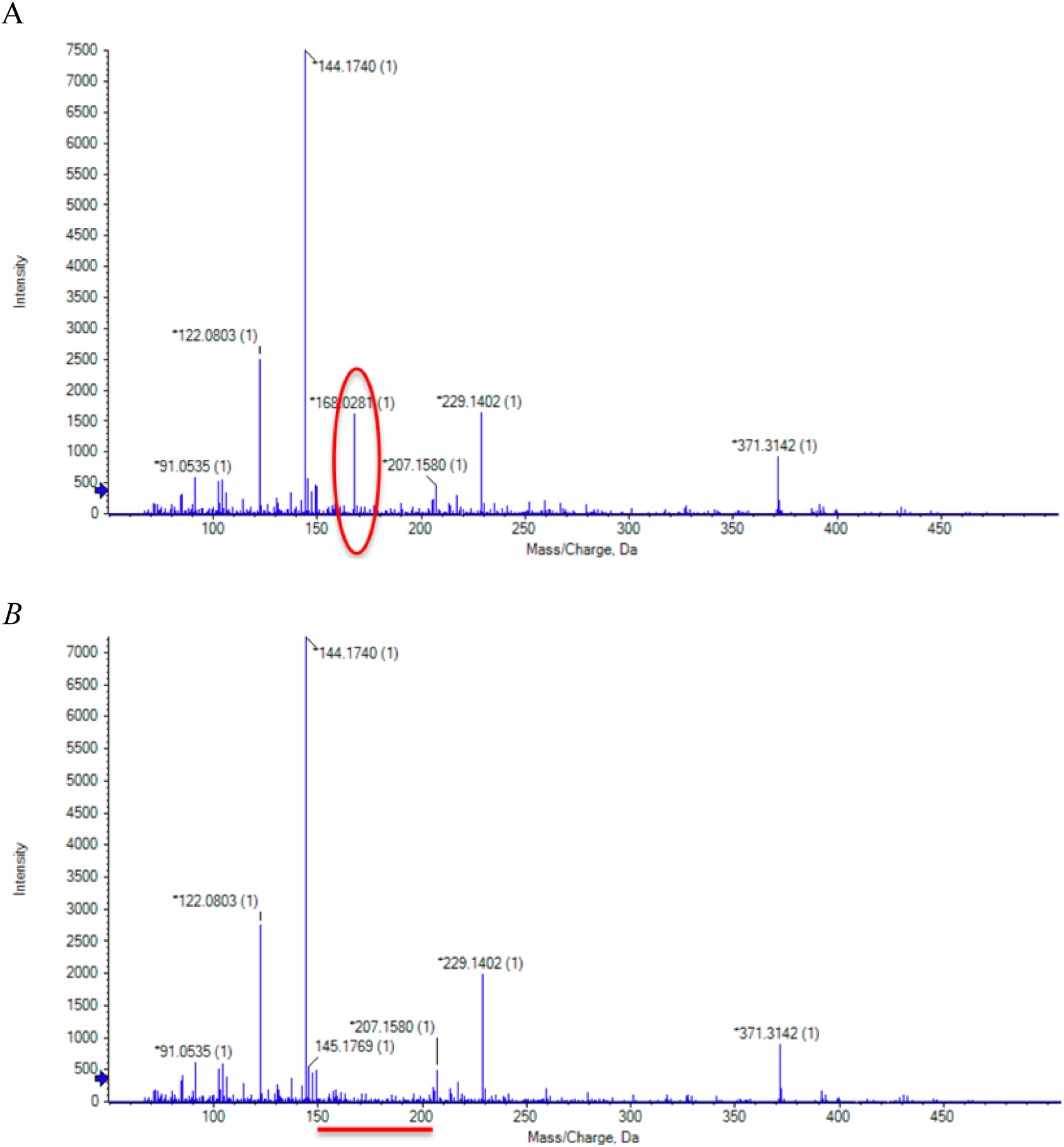
Detection of DPA by UHPLC-MS. Chromatogram depicting the detection of DPA (MW = 167.1 g/mol). (**A**) DPA (red circle) was detected in the spent germination buffer (50 mM Tris-HCl, 10 mM NaCl) from wild type *B. anthracis* spores. (B) DPA was not detected in the spent germination buffer from spores of *B. anthracis ΔspoVFAB.* Other DPA-less spores showed similar patterns and were omitted for clarity.

**Figure S4.**
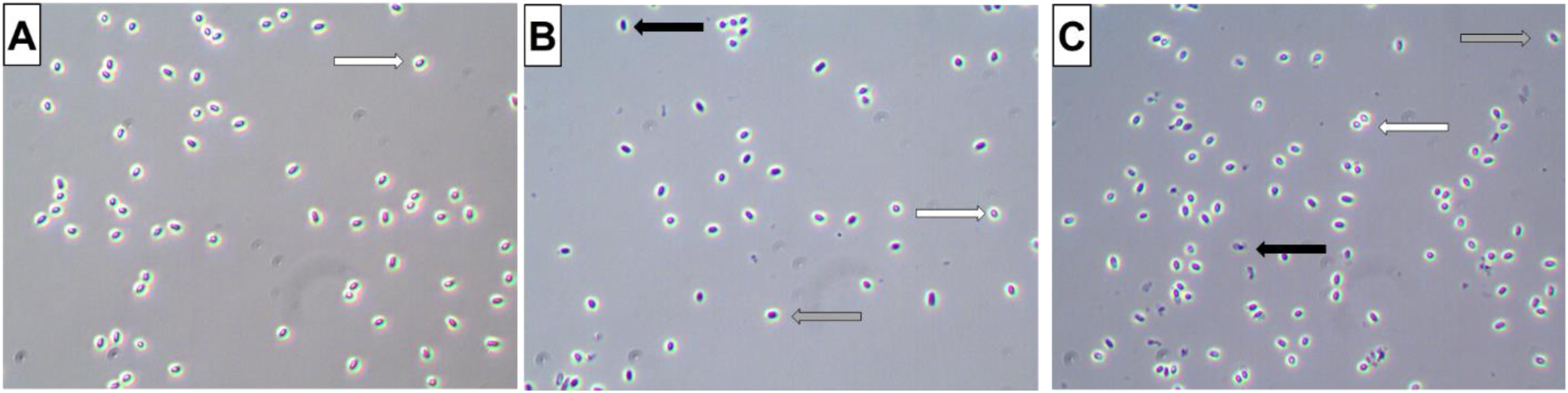
Phase contrast images of *B. anthracis* spores. (**A**) Only phase bright particles are detected in samples of wild type *B. anthracis* spores. (**B**) Samples of *B. anthracis ΔspoVFA* mutant spores contain phase bright (white arrows), phase gray (gray arrows), and phase dark (black arrows) particles. (**C**) Samples of *B. anthracis ΔsleBΔspoVFA* spores do not show an increase in the number of phase bright spores. Other DPA-less spores showed similar patterns and were omitted for clarity.

**Table S1.**
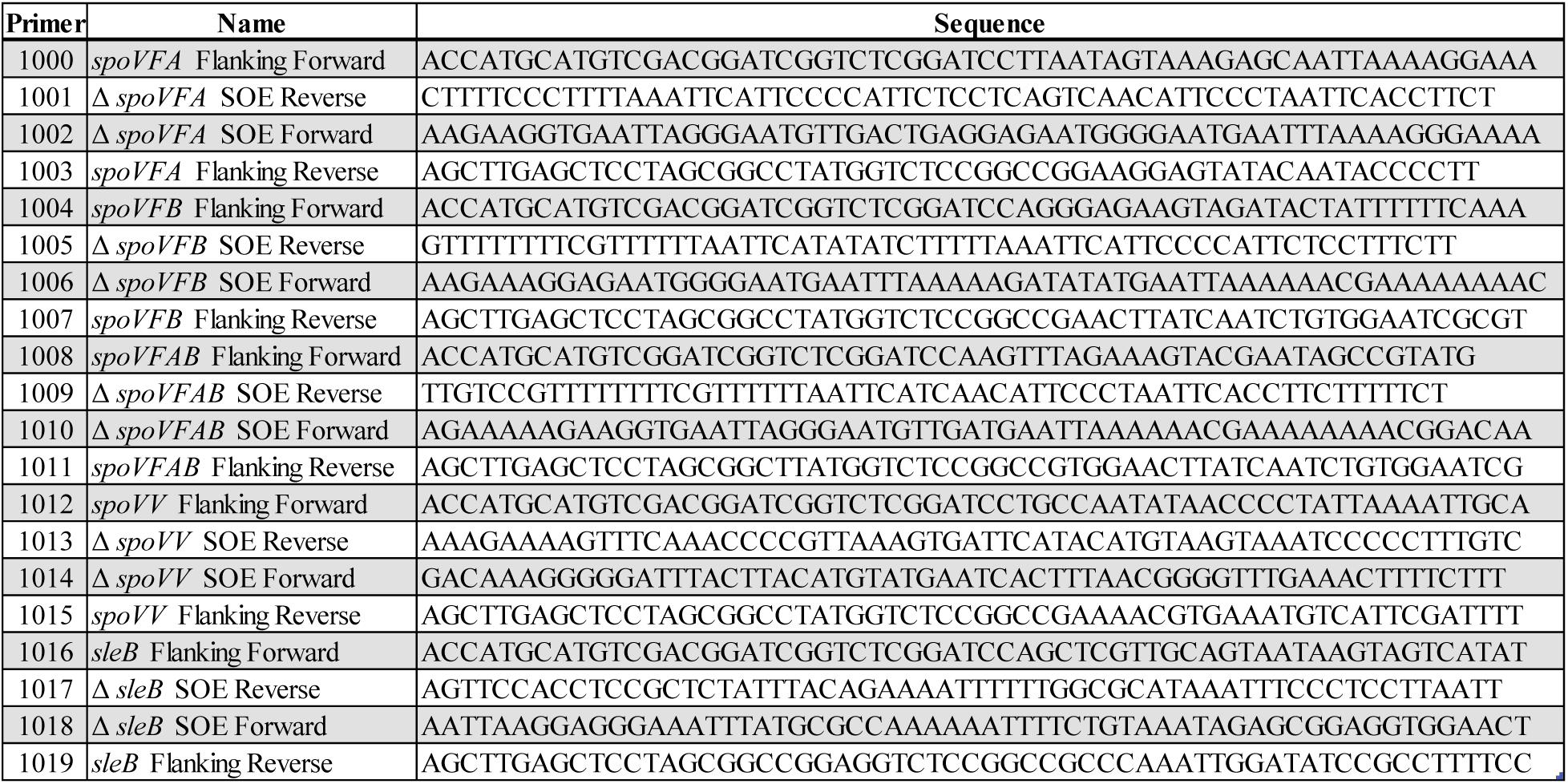
Primers used for knockout studies.

**Table S2.** Plasmids used in this project.

| Table S2. Plasmids used in this project |  |  |
| --- | --- | --- |
| Plasmid | Description | Source |
| pCR Blunt II-TOPO | Highly efficient cloning of blunt-ended PCR products; Km<sup>r</sup>, Zc<sup>r</sup> | Invitrogen |
| pRP1028 | Temp<sup>S</sup> allelic-exchange vector; Sp<sup>r</sup> | Dr. Roger Plaut (USDA) |
| pRP1099 | Facilitator plasmid; Km<sup>r</sup> | Dr. Roger Plaut (USDA) |
| pTOPO-ΔspoVFA | Cloning vector containing spoVFA SOE | This work |
| pTOPO-ΔspoVFB | Cloning vector containing spoVFB SOE | This work |
| pTOPO-ΔspoVFAB | Cloning vector containing spoVFAB SOE | This work |
| pTOPO-ΔspoVV | Cloning vector containing spoVV SOE | This work |
| pTOPO-ΔsleB | Cloning vector containing sleB SOE | This work |
| pRP-ΔspoVFA | Knockout vector containing spoVFA SOE | This work |
| pRP-ΔspoVFB | Knockout vector containing spoVFB SOE | This work |
| pRP-ΔspoVFAB | Knockout vector containing spoVFAB SOE | This work |
| pRP-ΔspoVV | Knockout vector containing spoVV SOE | This work |
| pRP-ΔsleB | Knockout vector containing sleB SOE | This work |

**Table S3.**
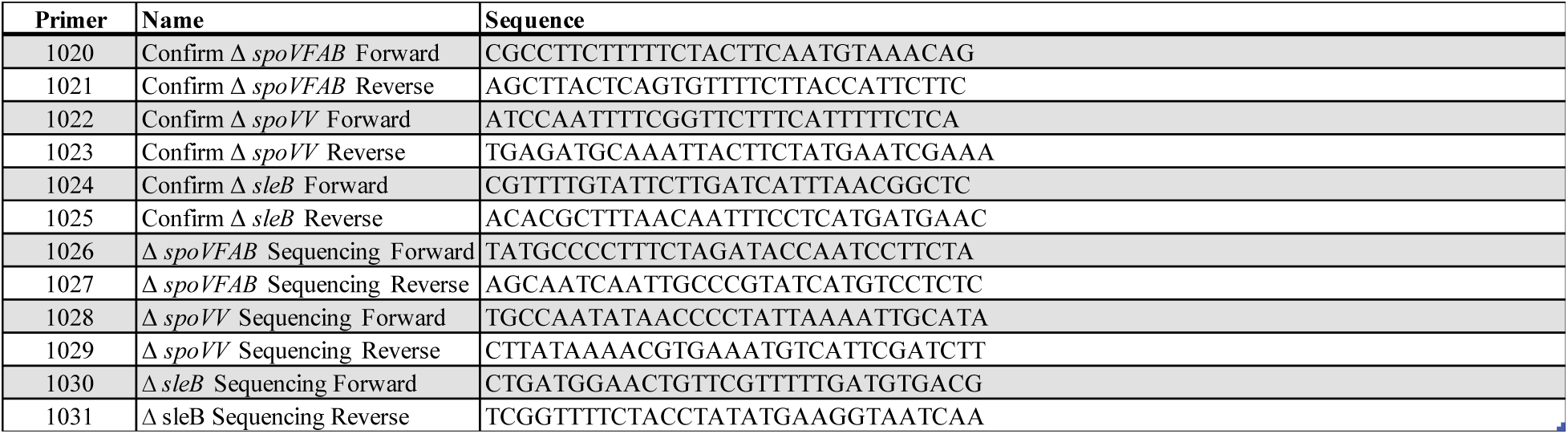
Primers used for *B. anthracis spoVFAB, spoVV,* and *sleB* sequencing.

**Table S4.** DPA released by wild-type and DPA-less *B. anthracis* spores.

| Table S4. DPA released by wild-type and DPA-less *B. anthracis* spores |  |  |
| --- | --- | --- |
| Spore Type | DPA concentration (mmol) | SD |
| Wild type | 35.7 | 4.8 |
| $\Delta spoVFA$ | 0 | 0 |
| $\Delta spoVFB$ | 0 | 0 |
| $\Delta spoVFAB$ | 0 | 0 |
| $\Delta spoVV$ | 0 | 0 |
| $\Delta sleB\Delta spoVFA$ | 0 | 0 |
| $\Delta sleB\Delta spoVFB$ | 0 | 0 |
| $\Delta sleB\Delta spoVFAB$ | 0 | 0 |
| $\Delta sleB\Delta spoVV$ | 0 | 0 |
| Suspensions of *B. anthracis* spores were diluted to OD<sub>580</sub>= 1.0 in DI-H<sub>2</sub>O and boiled for two hours to release DPA. The samples were mixed with a solution of TbPV, and fluorescence was measured. DPA concentrations were determined by interpolation of standard curves. Data are presented as mean ± SD from at least three independent measurements. | | |

